# Sensitivity Levels: Optimizing the Performance of Privacy Preserving DNA Alignment

**DOI:** 10.1101/292227

**Authors:** Maria Fernandes, Jérémie Decouchant, Marcus Völp, Francisco M Couto, Paulo Esteves-Veríssimo

## Abstract

The advent of high throughput next-generation sequencing (NGS) machines made DNA sequencing cheaper, but also put pressure on the genomic life-cycle, which includes aligning millions of short DNA sequences, called reads, to a reference genome. On the performance side, efficient algorithms have been developed, and parallelized on public clouds. On the privacy side, since genomic data are utterly sensitive, several cryptographic mechanisms have been proposed to align reads securely, with a lower performance than the former, which in turn are not secure. This manuscript proposes a novel contribution to improving the privacy × performance product in current genomic studies. Building on recent works that argue that genomics data needs to be × treated according to a threat-risk analysis, we introduce a multi-level sensitivity classification of genomic variations. Our classification prevents the amplification of possible privacy attacks, thanks to promoting and partitioning mechanisms among sensitivity levels. Thanks to this classification, reads can be aligned, stored, and later accessed, using different security levels. We then extend a recent filter, which detects the reads that carry sensitive information, to classify reads into sensitivity levels. Finally, based on a review of the existing alignment methods, we show that adapting alignment algorithms to reads sensitivity allows high performance gains, whilst enforcing high privacy levels. Our results indicate that using sensitivity levels is feasible to optimize the performance of privacy preserving alignment, if one combines the advantages of private and public clouds.

## Background

DNA sequencing and the alignment of sequences are at the heart of applications such as precision medicine, forensics, medical or anthropological research [1], [2], to name a few. Next-generation sequencing (NGS) machines greatly improved the throughput of human DNA sequencing and thereby reduced the costs of DNA analysis to almost 1000$ per genome. Widely deployed sequencing machines (e.g., from Roche or Illumina) produce short sequences of nucleotides ranging from 30 to 100 base pairs (bp) with error rates around 0.1% [3].

A simplified genomic workflow is made of the following tasks. First, the raw sequences of nucleotides that sequencing machines produce, called reads, are aligned to a reference genome to obtain their location in the genome. Then, aligned reads are used as input of the variant calling step, which identifies the donor’s genomic variations (i.e., her/his genotype). Finally, subsequent research or medical applications compare a subject’s genotype with other genotypes, or simply study it at a given locus.

On the one hand, the privacy of human genomes is particularly sensitive to data leaks. For example, not only do they uniquely identify their owners, but they also reveal information about their close relatives. In addition, once a genome has been revealed, its privacy cannot be recovered, as a subject’s genome barely evolves during his life. As multiple studies have shown, anonymizing human genomes, or creating aggregates, does not fully protect the privacy of their donors. To name a few, published privacy attacks included re-identification attacks [4], [5], and disease revealing attacks [6]. It has also been shown that leaking raw reads can expose their donors to data identity leakage [7], [8]. Consequently, besides standard encryption-based solutions, several works have been published to protect the advanced uses of genomic data: masking information in aligned reads [9], creating privacy-preserving releases of aggregated data [10], classifying raw genomic data as sensitive or non-sensitive [11].

On the other hand, efficient workflows are required, since the decrease of the sequencing prices has led to larger data productions. The e-biobanking vision [12], [13] argued for distributed and high-performance environments to host genomic workflows. Takai-Igarashi et al. proposed a data sharing policy which provides privacy protection in a biobanking environment [14]. Some more research have been done in order to develop privacy-preserving environments for genomic data computations, which include, but are not limited to, performing Genome-Wide Association Studies without revealing private data [15], and the development of methods to share aggregated data while preserving privacy [16]. Such works have been enabling the open science paradigm in the biological sciences, and progress in sharing of patient-derived health data has been more moderate [17]. However, moving genomic data to the clouds, among other possibilities, raises privacy challenges. Recently, several ecosystems addressing some of these challenges appeared. NGS-Logistics [18] allows the research community to access and analyze rare genomic variants while preserving the privacy of donors and the confidentiality of data. In particular, it relies on different levels of access rights for better protection of the data. Global Alliance [19] also proposed an ecosystem that allows remote accesses to databases located all around the world.

To summarize the current situation, previously published privacy attacks alerted the research community about the need to protect the genomic data, and the fact that many solutions currently in use may not protect data sufficiently, whereas other, stronger protection mechanisms are not efficient enough. There is therefore an urgent need to find the right balance between data protection and performance.

In this work, we focus on protecting sensitive genomic data both as soon as it is produced by an NGS machine – before the genomic variations they contain have been determined – and throughout the alignment step. As previous works [20],[18] argued, classifying genomic data as either sensitive, or not sensitive at all, is not sufficient. Building on these works, we remarked that a finer grained sensitivity classification of raw reads in combination with alignment algorithms which have different privacy guarantees and efficiencies, has the potential to improve the performance and overall privacy of the analysis workflow.

Our manuscript makes the following contributions:

1. We present a classification of raw reads into sensitivity levels, based on qualitative and quantitative characteristics of genomic variations. These sensitivity levels are then further partitioned in such a way that an adversary observing a part of the reads of a given sensitivity level, thanks to a successful attack, is not able to infer any more sensitive information from it. We disconnect sensitivity levels based on the Linkage Disequilibrium of genomic variations, and based on MaCH [21], a state-of-the-art haplotype inference software.
2. Building on previous work, namely [11], we propose to use a detection method based on Bloom Filters (BFs) to efficiently classify raw reads into partitions of sensitivity levels. In particular, we show how to preserve the disconnection property of sensitivity levels when Bloom filters produce false positives among the same or different sensitivity level.
3. We show that given a realistic heterogeneous and distributed environment, one can rely on the diversity of the existing alignment procedures to optimize the privacy *×* performance product of the read alignment step.

Whenever a public cloud is available, and is at least as powerful as the private infrastructure, our performance evaluation shows that our approach requires 0.29 CPU seconds and only 1.6 KB of data transfer to securely align a single read. Compared to a public cloud only approach, this represents a 10^6^-fold reduction of the computing time and 10^7^-fold reduction of the amount of data transferred to the cloud.

### Related Works

#### Privacy attacks on genomic data

DNA analysis is used in several fields, among them medicine, research, and forensics [1], [2]. After the sequencing process, the produced reads are typically used either in a *de novo* assembly, or aligned to a reference genome. Besides the wide use of genomic data on diverse fields, privacy attacks [8], [22] and the use of clouds environments for biomedical data analysis [23], [24], [25], [26] have raised concerns about the data security on the alignment step. Melissa Gymrek et al. [5] performed a re-identification attack identifying 131 of the anonymized genomes of the 1000 Genomes Project. Nyholt et al. [6] reconstructed sufficient evidence to deduce the likelihood of Prof. Watson getting Alzheimer’s disease, although he deliberately removed the APOE’s gene before publishing his genome. Homer et al. showed that is possible to determine if a subject contributed with his DNA to a mixture using their DNA profiles [4]. Goodrish described a mastermind mind attack where querying successively encrypted data may allow to discover private data [27]. Kidd et all developed a method that uses a high probability SNPs panel to determine the ethnicity of subjects [28]. Some studies contributed to the definition of trail attack, which consists on the use of unique distinguish features collected from different institutions to match DNA samples and their donors [29], [30], [31].

#### Privacy preserving mechanisms for genomic data

The reported privacy attacks, which we summarize on the previous section, made genomic privacy a priority challenge of the biomedical community [32], [33], [34]. In order to address this challenge, the biomedical community invested on strategies to protect genomic data privacy and defined three categories of protection: data de-identification [35], [31], data augmentation [36], [30], and protection recurring to cryptographic methods [27], [37].

**Data de-identification methods:** These methods remove the personal identifiers, such as names, and social security numbers, which initially connects the genomic data to its owner. Neubauer et Heurix [38] proposed a methodology to protects medical data through pseudonymization. However, alone these methods are not sufficient to provide privacy protection of genomic data, since they do not complete solve the re-identification problems [35], [31], [39].

**Data augmentation methods:** This category of data protection uses generalization to achieve privacy protection. The basis of generalization is to make all the records indistinguishable from all the other shared records. This methods achieve better protection than the previous category, however some data applications are lost.

**Protection recurring to cryptographic methods:** The cryptographic methods use privacy-preserving querying methods designed for sequenced data which allows their utility [27], [37], without accessing to the original data. However, the protection provided by methods that rely only on encryption mechanisms can be lost in a shorter time than genomic data privacy requires [40], which makes these methods not efficient.

#### Existing reads alignment methods

In this manuscript, we consider the following three main types of alignment algorithms, which we include later in our discussion of privacy protection per sensitivity level.

**Distributed plaintext alignment:** This category contains fast algorithms, which are usually used in clouds to enable parallel study of large amounts of data. However, these algorithms do not consider any privacy-preserving mechanisms. Some examples of those algorithms are CloudBurst [41] and DistMap [42] which perform plaintext alignments in public clouds (either with or without encrypting the data transferred).

**Proven-secure distributed alignment:** The main goal of this second category is privacy. Algorithms in this category are secure – they do not present risks of data leakage – but they achieve this by sacrificing performance, and thus they are much slower than algorithms of the previous category. Garbled circuits alignment [43] and homomorphic encryption schemes [44] have been used to perform operations on encrypted reads.

**Non-proven-secure distributed alignment:** Considering the drawbacks of the previous two categories, in the last years researchers have been searching for intermediate solutions. These solutions try to combine high performance and security. Chen et al. [45] proposed a seed-and-extend alignment implementation in hybrid clouds environment. In their approach privacy is maintained thanks to the transmission of reads hashes to the public cloud. All the privacy sensitive steps are performed in a private cloud. A different approach is followed by Balaur [46], which uses Locally Sensitive Hashing (LSH), secure k-mer voting, and a MinHash algorithm to align reads. We selected Chen et al.’s method for our experiments, as it is lighter in terms of memory consumption.

#### Detection and sharing of sensitive raw information

Ayday et al. [9] proposed the storage of encrypted short reads in a biobank. This solution enables classified people (e.g., data analyst in a hospital) to retrieve a subset of the reads to perform genetic tests without revealing their nature to the biobank. In the described solution, the biobank can mask some regions of the short reads to better protect them. For example, regions outside the request range, or regions for which the patient does not give consent to access. Cogo et al. [11] describe a filtering approach that based on a knowledge databases and supported by Bloom filters is able to work on 30 bp sequences, but does not support privacy levels. Bloom filters [47] are a data structure that is used to test whether a given element is part of a predefined set. Schnell et al. [48] proposed a privacy-preserving record linkage method using Bloom filters to compute the similarity of encrypted identifiers.

## Methods

### Data, system and threat model

**Data:** We build the sensitivity levels based on the set of genomes available from the 1000 Genomes Project [49]. We also recombine genomic variations with the reference genome GRCh38.p11 to create possible genomic sequences that carry genomic variations. Such sensitive sequences are used to initialize read filters [11], and classify reads into sensitivity levels.

**System model:** We consider a biocenter whose task is to generate reads from an individual, and to align those reads to a reference genome in a privacy-preserving manner. To do so, the biocenter can rely on a private cloud, and public clouds. We assume that the sequencing machine and the private cloud are secure. However, we consider a private cloud expensive to maintain, which encourages the use of public clouds, even though we assume that the user does not have a complete control over its own data (i.e., which machines are used, etc.) in public clouds. Finally, we consider that all parties can rely on encrypted communication channels, if needed, and that they use unbreakable cryptographic procedures.

**Threat model:** We assume an honest-but-curious adversary, which tries to observe and learn sensitive genomic information during the alignment of reads. In particular, the adversary is able to observe raw reads in the public cloud if they are used in plaintext alignment algorithms. We also assume that the adversary has access to a reference genome, is able to align raw reads to obtain the biological insights they contain, and has access to the statistical relationships between genomic variations. Obtaining such refined data can then potentially enable existing privacy attacks during future uses of data (e.g., if allele frequencies in a case population are publicly released). To limit the risk that an adversary obtains sensitive information while obtaining high performance using cleartext alignment, we use a risk-threat approach to align and protect the reads.

**Amplification attack:** In addition, as perfect security does not exist, in case a successful attack happens, where an adversary would be able to observe raw reads, we aim at preventing this attack from being extended to data that could not be observed directly during the attack (e.g., because it is more protected, or used in a different location). We call this an amplification attack. We use Linkage Disequilibrium (LD) measurements, and MaCH [21], a state-of-the-art haplotype inference software, to make sure that an adversary is never able to execute such amplification attacks.

### Sensitivity levels

In this section we describe the first step of our classification method that consists in the creation of the different sensitivity levels. We explain how to create the different levels in such a way that privacy leak amplification is not possible, namely using cloud diversity and promotions of genomic variations across levels.

### Qualitative and quantitative sensitivity levels

In this manuscript we propose the creation of sensitivity levels that allow the differentiation of genomic data and basically can be a combination of two different methods: manually declaration (qualitative), or based on frequencies in a reference population (quantitative). In the first case, the researcher decides the sensitivity levels based on how he perceives the sensitivity of the information a variation reveals. Sensitive levels based on frequencies, as we propose in this manuscript, are built on the fact that a rare disease/genetic variation should be considered more sensitive than a common one, since they concern a smallest subset of the population. In particular, alleles whose frequencies are lower than 0.05 should always be considered highly sensitive since they can conduct to a restricted group of individuals [50].

Figure 1a shows, as an example, a distribution of genomic variations in three sensitivity levels, based on the frequency of the alleles in the genome. In this figure, level 1 contains the alleles whose frequency goes up to 0.05, the level 2 is composed of the alleles whose frequency is comprised between 0.05 and 0.2, and the level 3 holds the remaining more common alleles. Later in the results section, we discuss the distribution of the alleles among different possible sensitivity levels. Differently, Figure 1b presents three sensitivity levels based on information coded in the genome and their severity when leaked. For level 1 we consider information such as disease genes, ethnicity related variants and other regions that can lead to individual’s re-identification.

**Fig. 1.**
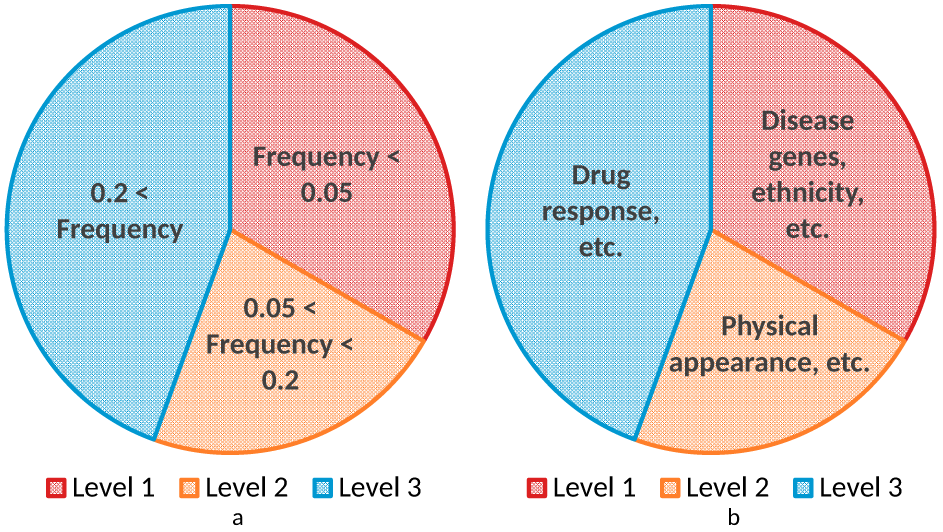
Initial sensitivity levels. (a) Alleles frequency-based sensitivity levels (quantitative classification). (b) Manual declaration-based sensitivity levels (qualitative classification).

### Preventing direct LD inference

Linkage disequilibrium (LD) describes the non-random transmission of genomic variations [51]. These non-random associations of genomic variations have been used in privacy attacks [8].

We computed the LD between the genomic variations in the human genome using a maximum range of 1000 bases between two variations. We show the results in Figure 2a, where as example we show variation 1 linked with variation 3, which is linked to variation 6. In this case we have direct inference connections, which should be avoided, between all the three sensitivity levels. Indeed, if an adversary obtains the variation 3, he will be able to identify the variations 1 and 6. Then we have two more cases of linkage between variations, variation 2 is linked to variation 5, and variation 4 is connected with variations 7 and 8. This range considers that LD relations can extend to few kilobases (Kb), and sporadically up to 100 kb [52].

**Fig. 2.**
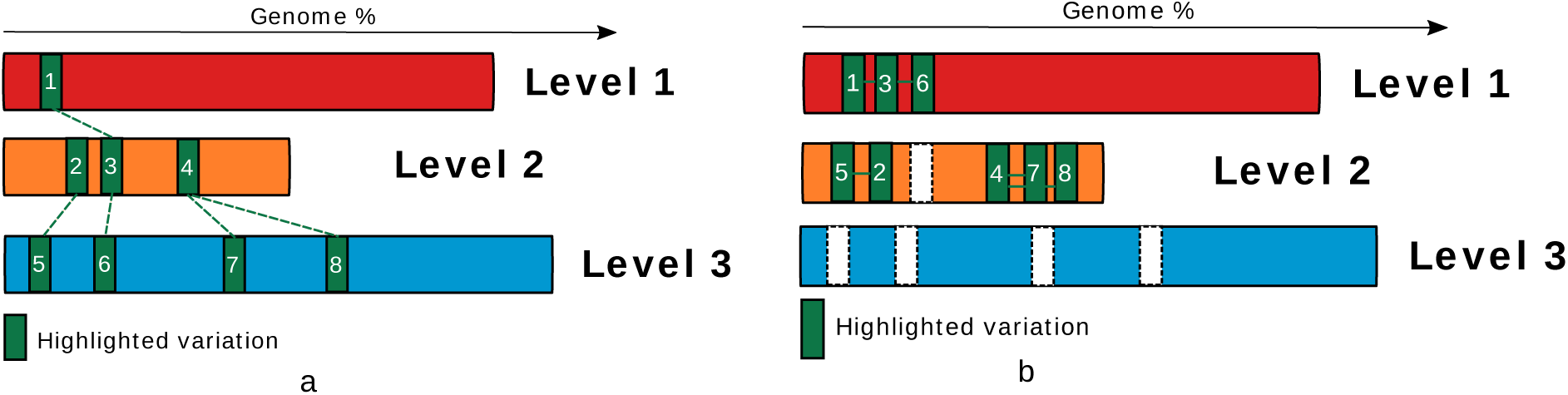
Allele promotion based on linkage disequilibrium (LD). (a) Variations in LD are identified on the initial layers. (b) Alleles levels refinement after LD-based promotions.

To avoid a similar attack as described above, after we have computed all sets of related genomic variations through LD, we promote the variations spread across different levels to the highest sensitivity level (Figure 2).

### Sensitivity level promotion through haplotypes inference

To ensure that sensitivity levels are completely disconnected, e.g., that there will remain no-linked single nucleotide polymorphisms (SNPs) across sensitivity levels, we use a haplotype inference software (e.g., MaCH [21]) which is able to infer missing SNPs for an individual of a reference population. From a sensitive level, we compute the higher sensitive genomic variations that can be inferred with an *r*^2^ value higher than 0.3 (value recommended by MaCH’s authors). The software receives as input two sets: (i) a list of the biomarkers and the SNPs information of a reference population, and (ii) a list of the biomarkers and the SNPs information the adversary can observe. To get the disconnected sensitivity levels we perform the following steps for each level:

**Step 1. Create the input files to run MaCH.** We start by creating two pairs of files based on subsets of 20,000 SNPs from Chromosome 1, available in the 1000 Genomes Project. We believe this number to be high enough to be representative, since the correlation of SNPs decreases with their distance. The first pair of files (i.e., the .snps and .haplos files) details the set of complete genotypes the adversary uses as reference. The second set of files (i.e., the .dat and .ped files) details the genomic variations the adversary is able to observe at a given sensitivity level (through an hypothetic attack) and masks the more sensitive SNPs which would be located in a secure environment.

**Step 2. Run MaCH for inference.** We run MaCH with the input files, and obtain a list of SNPs that can be inferred by an adversary with the provided sets, i.e., masked SNPs that MaCH is able to discover with good accuracy. We focus on the inference of more sensitive SNPs, thus we ignore those that are inferred and belong to the same sensitivity level. This step assesses the information that can be inferred in case of information leakage from a sensitivity level.

**Step 3. Find the SNPs correlated with the ones found by MaCH.** From the inferred SNPs, we compute the top 10 SNPs related to each inferred SNP that the adversary can observe. MaCH only provides the list of inferred ones, not the relations between them, thus we computed the linkage disequilibrium between the SNPs set input as observed by the adversary and the set of inferred SNPs. We remove 10 SNPs at each iteration to speed up our experiments. Using lower values would reduce the number of promotions, but would increase the number of iterations, which would be more time consuming.

**Step 4. Remove the correlated SNPs from the input files.** We remove the top 10 related SNPs from the initial adversary set, since they allow the inference of more sensitive SNPs. From these newly obtained input files, we iterate again, until no more inferences are possible, or their number stabilizes.

Figure 3 provides an illustration of the inference iteration process using MaCH on the example previously introduced in Figure 2a, after alleles in strong direct LD have been promoted. In this example, MaCH observes the genomic variations contained in sensitivity level 3 (created during step 1.), and tries to infer more sensitive genomic variations (step 2.). After inference, MaCH infers that the genomic variation numbered 4, which is in level 2, can be inferred with good accuracy from those in level 3. Our code would then identify that the genomic variations numbered 7 and 8 in level 3 are strongly associated with the one numbered 4 (Step 3.). Those two genomic variations would then be promoted to level 2. (Step 4.), before iterating the inference process.

**Fig. 3.**
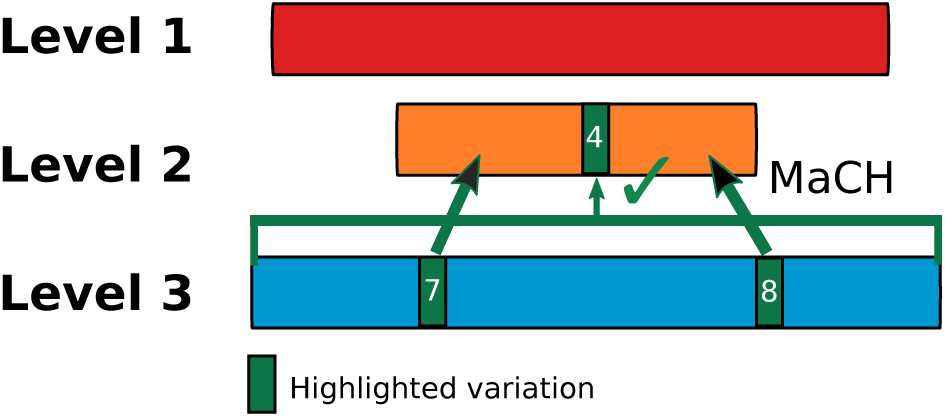
Information stored in a single cloud. Allele promotions based on inference performed by MaCH.

### Classifying sensitive reads and adapted treatment

In this section, we first recall how sensitive reads (i.e., reads that carry sensitive private personal information (PPI)) can be detected thanks to the filtering method proposed by Cogo et al. [11]. We then explain how to extend that method to classify reads into several sensitivity levels (i.e., not just based on a binary answer) depending on the magnitude of the impending privacy risk – the probability of an adversary extracting PPI, and the resulting negative impact. Finally, we show how to solve possible detection conflicts when using several filters.

#### Short read filter

We briefly introduce the short reads filtering method [11] and how we can use it to classify reads into a scale of sensitivity levels. Cogo’s read filter is implemented using a Bloom filter, which is a high throughput data structure that can produce false positives, but never false negatives. In addition, the filter is not a bottleneck when used in combination with sequencing machines, as its throughput is always at least 40 times faster than the throughput of current NGS machines, and it is parallelizable.

The filter is initialized from a database of reads known to carry sensitive information. This sensitive information includes, but is not limited to, all existing data that have been used in the literature that describe attacks to re-identify subjects of experiments. Such attacks have been based on three kinds of sensitive sequences: (i) genomic variations, (ii) disease genes and (iii) short tandem repeats (STRs) – a known small string that appears several times contiguously in a subject’s DNA, and whose repetition numbers vary among a population.

The filtering process is initialized thanks to three offline steps: (i) build a dictionary of sensitive levels; (ii) refine the sensitive levels; and (iii) initialize the Bloom filter from the dictionary. Finally, once the Bloom filters has been loaded into memory, a further online step filters raw reads into the levels, and outputs them.

#### Classification into sensitivity levels

We use one short read filter per sensitivity level that we have previously identified, to prevent amplification attacks. To build the dictionary of sensitive levels, we collect all genomic variations of a subgroup, and create all possible 30 bp sensitive sequences from the genomic variations and the reference genome, and build the associated dictionary of sensitive sequences. Each dictionary is then inserted into a Bloom filter, which is then ready to filter sequences.

Figure 4 illustrates the filtering approach we use, where the goal is to classify and group reads separately, according to their sensitivity level. The filtering approach has three steps: sequencing, sensitive-aware filtering, and conflict management, which is needed in case a read matches in several filters. In the sequencing step, the DNA sample is translated into reads by the NGS machine. The reads are then given as input to the sensitivity-aware filtering step, made of Bloom filters initialized with several disconnected groups of genomic variations. Finally, the last step consists of collecting the outputs of the Bloom filters, and deciding the sensitivity level of a read, and the subgroup it should be part of. Depending on the sensitivity of a read, it becomes then possible to adapt its storage, computation, and use according to the security required. As represented in Figure 4, storage costs and access limitations per read tend to increase with the sensitivity of read, while alignment performance tends to decrease. Specific numbers depend on the available infrastructures, and on design choices.

**Fig. 4.**
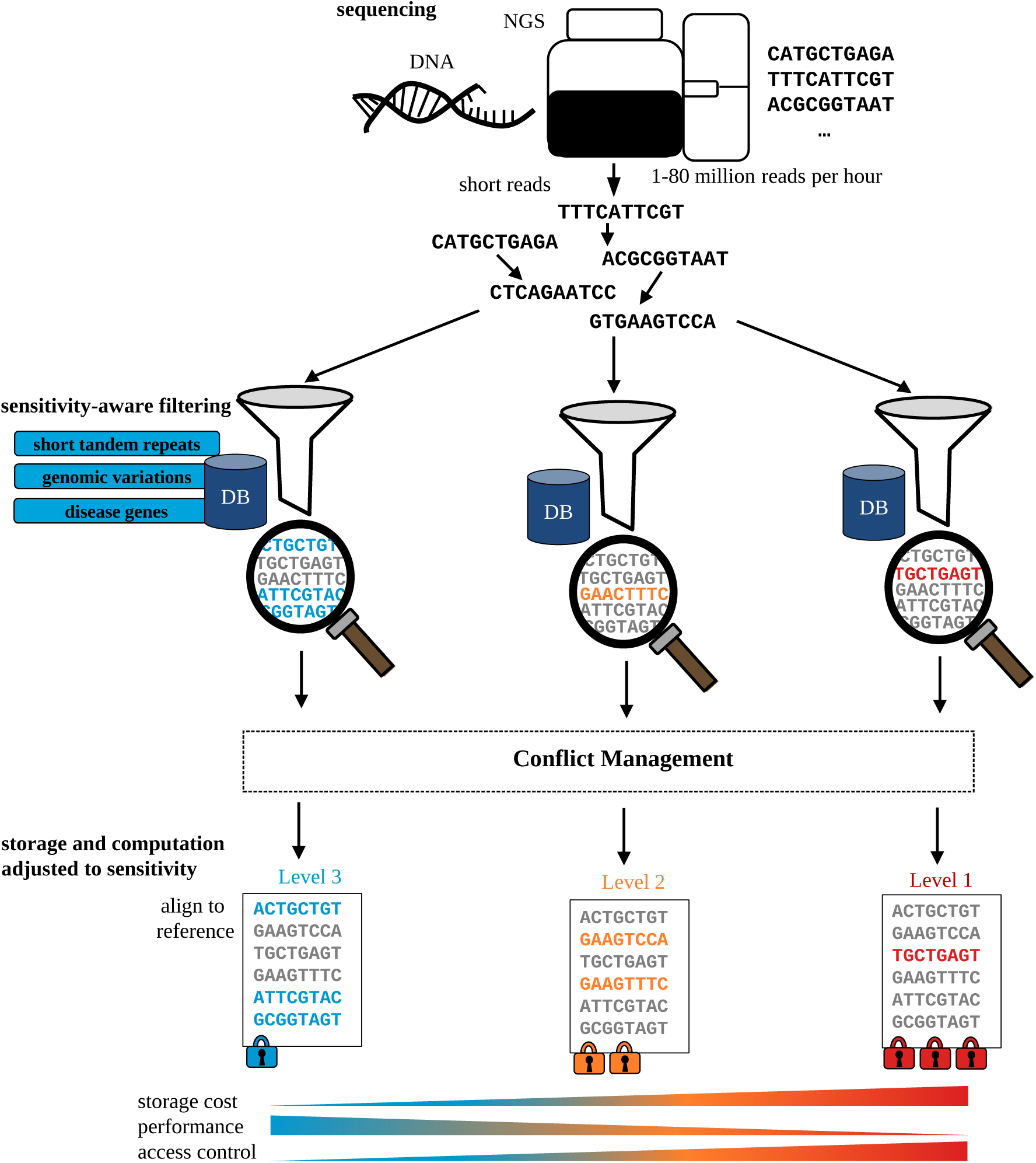
Filtering of reads based on sensitivity levels. After creation of the sensitivity levels we use Bloom filters to classify reads into the created levels. After the classification we can adjust the storage and algorithms, and control the access per level.

#### Filters conflict management

Bloom filters may produce false positives, which in our case may cause privacy leaks if they are not correctly managed. Handling false positives is part of the conflict management step represented on Figure 4, and works as follows. If a read matches in several sensitivity level filters, we set its sensitivity to the highest existing sensitivity level. This is necessary since if a read belongs in a lower sensitivity set, and is also detected as belonging in a higher sensitivity level, due to a false positive, then inserting it in the higher sensitivity set may enable an amplification attack. More precisely, the union of the higher sensitivity set and this read’s genomic information have not been controlled when sensitivity levels were initially disconnected (i.e., using MaCH). The number of these few reads can be controlled by the false positive parameters of Bloom filters.

### Read alignment: Performance × Privacy product optimization

We finally adapt the read alignment step to the detected sensitivity of the reads. We show that it is possible to optimize the performance x privacy product of read alignment by adapting the alignment algorithm to the detected sensitivity of a read.

We study several scenarios, and measure the performance improvement against standard alignment mechanisms.

*Scenario 1: public clouds only*. Read alignment is performed in a non-secure environment where unencrypted computations and communications may be intercepted by an adversary. In this case, sensitive reads have to be aligned with believed or proven secure algorithms. However, more efficient algorithms can be used on low sensitivity reads.

*Scenario 2: sensitivity-adapted private and public clouds alignment.* High-sensitivity reads are aligned in a private cloud, while low-sensitivity reads can be aligned in public clouds. In practice, biocenters have computing resources, and use them for their most sensitive computations. We show that we can extend those computing resources, by a secure usage of public clouds.

## Results

For our performance evaluation experiments, we rely on both a private cloud, and public clouds under different scenarios. To align reads using the sensitivity levels, our approach always aligns the most sensitive reads in the private cloud (using CloudBurst [41]), and the least sensitive reads in the public cloud (using CloudBurst [41]). The remaining reads are either aligned in the private cloud (using CloudBurst [41]) or in the public cloud (using Chen et al.’s protocol [45]), depending on which cloud finishes first.

### Sensitivity levels statistics

We studied the average proportion of a subject’s SNPs in each sensitivity level before and after SNPs promotion through haplotype inference. Figure 5a represents the proportion of genomic variations of a subject in each sensitivity level before the promotions. Level 1 contains a minority of alleles (3%), level 2 contains only 2% of the alleles, and the remaining 95% lies in level 3.

**Fig. 5.**
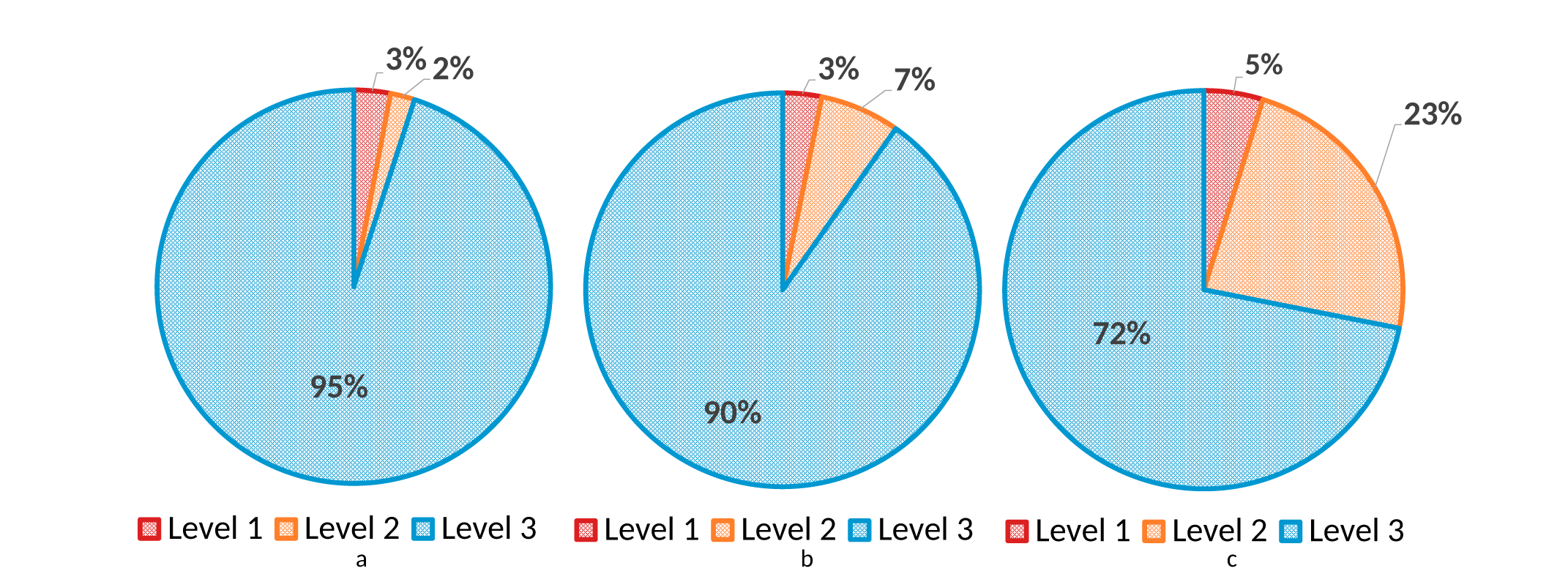
Sensitivity levels evolution through promotions process. Red color represents the most sensitivity level (1), orange the level 2, and blue the least sensitive level (3). (a) Represents the proportion of an individual genomic variations (GVs) per sensitivity level. (b) Proportion of an individual GVs per sensitivity level after promotions. (c) Proportion of 100-bases reads per sensitivity level.

The genomic variations promotion slightly change the distribution among the sensitivity levels, as Figure 5b shows. In this case level 1 is the smallest one with 3%, level 2 slightly increases with now 7% of the alleles, and the last, level 3, contains 90% of the alleles.

### SNPs promotion across sensitivity levels

Overall, after one iteration, we promoted 1.6% of the SNPs of level 1.0 and 18% of the SNPs of level 0.2. Overall, we promoted 1.5% of all SNPs from one level to a more sensitive one. We summarize the proportion of inferred SNPs per sensitivity level after various rounds of promotion iterations, in Figure 6. The promotions are made using the method described in section. After one inference iteration with MaCH, very few genomic variations could still be inferred (e.g., less than 5 SNPS with level 3). We believe that these inferences are due to the limited number of genomes used in the 1000 Genomes project, and to specific individuals who had unique combinations of statistically unlinked SNPs (since they can still be inferred after more iterations). We are confident that inferring those SNPs in a larger population would not be possible because the number of unique combinations of several SNPs would be rarer.

**Fig. 6.**
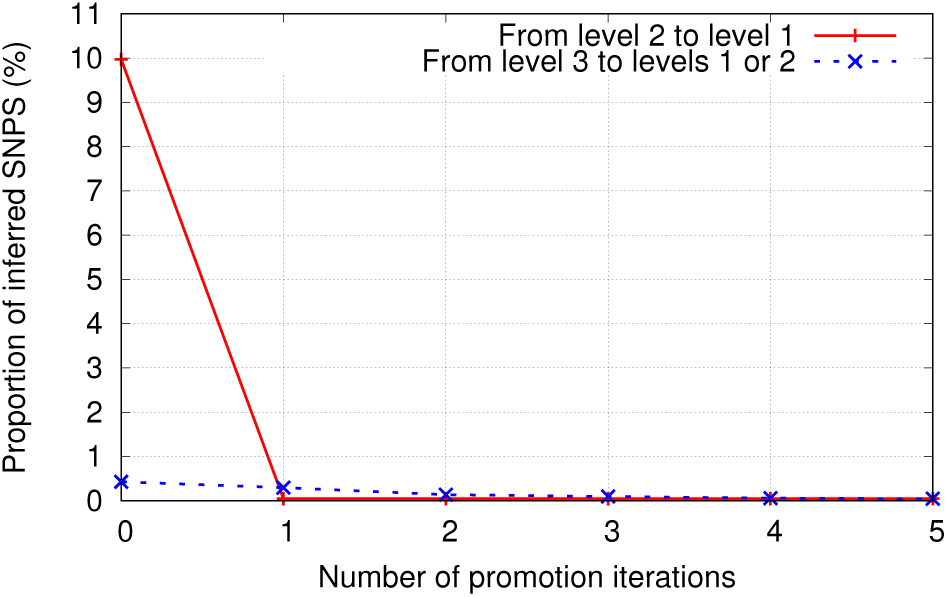
Number of inferred SNPs per inference and promotion. The red line represents the promotions from level 2 to level 1 and the blue line represents the promotions from level 3 to levels 1 and 2.

### Reads classification on sensitivity levels

Figure 5c shows the percentage of 100-bases reads in each sensitivity level. This distribution is somewhat different from the distribution of the genomic variations. Considering reads, level 1 contains only 5% of the reads maintaining the most sensitive level with the lowest amount of information. Level 2 contains 23% of the reads, and the remaining 72% of the reads are classified into level 3. Level 3 continues to hold the majority of the information which support the performance and privacy optimization we discuss in next section.

### Performance × Privacy product optimization

Aligning reads in a cloud implies assuming that the cloud provider is trustful, and that the cloud will not be attacked. We do not make these risky assumptions, and therefore rely on the following three categories of alignment algorithms, which we previously introduced in Section, to optimize the performance × privacy product using sensitivity levels. Category (i) – *non-secure but fast algorithms:* we use Cloudburst [41], which is an unprotected method, and requires 0.4 CPU seconds, respectively 0.41 CPU seconds if reads are encrypted for the transfer to the cloud server. Category (ii) – *secure but slow cryptographic algorithms:* we use a homomorphic encryption based approach [53], which requires 22 CPU days. Category (iii) – *algorithms providing an intermediate level of protection:* we use a protocol based on hashing k-mers presented in [45], which is much more efficient, requiring only 1.3 CPU seconds. However, it may leak information about equal k-mers and has not been formally proven theoretically secure.

Table I summarizes our analysis of the privacy level, computation cost (CPU hours) and communication (bytes) cost of aligning a single 100 base-pairs read to the full human genome. The values in the table are computed for an alignment using a single core.

**TABLE I.** Summary of the Privacy, Performance and Communication Overheads of the Alignment Algorithms We Consider.

| Method | Privacy<br>(Sec. ) | Computation<br>(CPU time) | Communication<br>volume |
| --- | --- | --- | --- |
| Homom. encr.<br>[53] | Very high | 22.08 days | $3.75 \times 10^8$ KB |
| Hashed k-mers<br>[45] | High | 1.3 sec. | 5.22 KB |
| Cloudburst<br>[41] | Low | 0.41 sec. | 2.3 KB |

Table II lists situations with different relative proportions of public cloud’s computing power over the private cloud computing’s power, while Table III presents the communication costs for the same proportions. For example, configuration 1/10 means that the public clouds are 10 times more powerful than the private cloud. Under each configuration we evaluate the performance of a read alignment for the three possible cases: (i) on the public cloud only (using 5PM [53]); (ii) on the private cloud only (using CloudBurst [41]); or (iii) on both the private cloud for sensitive reads and on the public clouds for non-sensitive reads (using 5PM [53]) on public clouds (using CloudBurst [41] on public clouds).

**TABLE II.** Computation Overhead of Existing Privacy-Preserving Approaches. We Compare Our Sensitivity-Adapted Alignment With the Previous Approaches and Depending on the Proportion of Public and Private Clouds Available.

| Proportion Pub./Pri. | Our approach | Previous approaches |  |  |
| --- | --- | --- | --- | --- |
| | | Pub [53]<br>($3 \times 10^8$s) | Priv [41]<br>(0.41s) | Pub. [53]/Pri. [41] with [11]<br>(0.29s) |
| 1/1 | 0.29s | $10^6$x | 1.39x | 1x (0.29s) |
| 2/1 | 0.097s | $10^6$x | 4.20x | 1.51x (0.11s) |
| 10/1 | 0.019s | $10^6$x | 20x | 5.85x (0.11s) |

**TABLE III.** Communication Overhead of Existing Privacy-Preserving Approaches. We Compare Our Sensitivity-Adapted Alignment With the Previous Approaches and Depending on the Proportion of Public and Private Clouds Available.

| Proportion Pub./Pri. | Our approach | Previous approaches |  |  |
| --- | --- | --- | --- | --- |
|  |  | Pub [53]<br>(16.8GB) | Priv [41]<br>(2.3KB) | Pub. [53]/Pri. [41] with [11]<br>(1.6KB) |
| 1/1 | 1.6KB | $10^7$x | 1.39x | 1x (1.6KB) |
| 2/1 | 0.55KB | $10^7$x | 4.20x | 1.51x (0.83KB) |
| 10/1 | 0.11KB | $10^7$x | 20x | 5.85x (0.65KB) |

Overall, we can draw the following conclusions about privacy-preserving alignment of a read: (i) it is not practical to rely only on a public cloud to align reads with cryptographically secure algorithms (3 × 10^8^ seconds per CPU per read); (ii) relying on a private cloud with cleartext alignment is the fastest solution (0.41 seconds per CPU), but it does not scale; (iii) by classifying reads as either sensitive or non-sensitive, performance can be improved whenever a public cloud, assumed to be as least as powerful as the private cloud, is available (starting at 0.29s with [11]); and (iv) our approach, relying on more than two sensitivity levels, further improves performance on hybrid clouds (down to 0.019s with a public cloud ten times more powerful than the private cloud). Similar conclusions can be taken in terms of memory consumption. To summarize our results, and compared to previous works, by using sensitivity levels to align reads, we remove computational tasks from the performance bottleneck of secure alignment, the private cloud, to execute them securely in public clouds.

## Discussion

In this manuscript we proposed a methodology to create sensitivity levels for raw reads and automatically detect the sensitivity level a read belongs to. Our methodology allows sensitive levels to be defined based on both qualitative and quantitative aspects. The levels declared qualitatively are based on the biological insights a sequence reveals, while the levels declared quantitatively are based on the frequency of variations in the general population. Since defining qualitative levels is subjective, we base our experiments on levels defined on quantitative aspects only, for simplicity.

We showed how to prevent leakage across levels due to haplotype inference (using LD relations), by promoting groups of linked sequences to the highest of their levels.

We then extended the filtering method proposed by Cogo et al. [11] to automatically classify reads into the multiple sensitivity levels (i.e., not just based on a binary answer). These results support the viability of a differentiated alignment for privacy × performance optimization.

Finally, we used our automatic classification mechanism to align reads. Our results show that classifying data into sensitivity levels improves performance during read alignment while it provides the adequate security to the processed data. We demonstrated our implementation to be more efficient than state of the art privacy preserving alignment methods, whenever a public cloud is available, with a computation time of at least 0.60 CPU seconds per 100 bases read and 2.9 KB of data to be transferred.

**Completeness of the databases:** Newly discovered genomic variations cannot be detected by the filter. However, updating the filter to include new sensitive genomic variations as they are discovered is straightforward. If the number of newly inserted sequences becomes too important, it may be necessary to create and initialize a larger filter, since inserting too many new elements in an existing Bloom filter increases its false positive rate. To answer this point, Cogo et al. [11] argue that new genomic variations are now rarely discovered, which limits the residual risk of not detecting a large number of yet unknown sensitive sequences. In addition, our approach limits the number of observable SNPs, and would decrease the potential for successful inference attacks.

**Extension to longer reads and higher error rates:** The filtering method we use classifies a full read into a sensitivity level. We showed that this approach is practical with short reads. However, a larger proportion of long reads would be classified as sensitive. Indeed, a longer read has a higher chance of containing more sensitive information than a short read. In addition, the probability that a long read contains several genomic variations increases. Extending this method to long reads would require modifications of the filtering algorithm. Another limitation of the filter is that it loses sensitivity and accuracy with reads sequenced with higher error rates. Even though possibilities exist to extend the filter to tolerate more errors, they are out of the scope of this manuscript.

**Cloud diversity to prevent inference:** Another solution to avoid amplification attacks could consist in splitting the input sets during MaCH experiments between different clouds. Precisely determining how to split a sensitivity level on different clouds could increase performance by reducing the number of promotions across sensitivity levels. Splitting each sensitivity level over several clouds would render amplification attacks even more difficult. In Figure 7, level 3 information is stored in two clouds (A and B). This division considers pairs of variation that are linked with each others, and ensure that they are separated and stored in distinct clouds. With this strategy, an adversary that obtains the content of one cloud (in the example, Cloud A) is not able to infer the higher level variant.

**Fig. 7.**
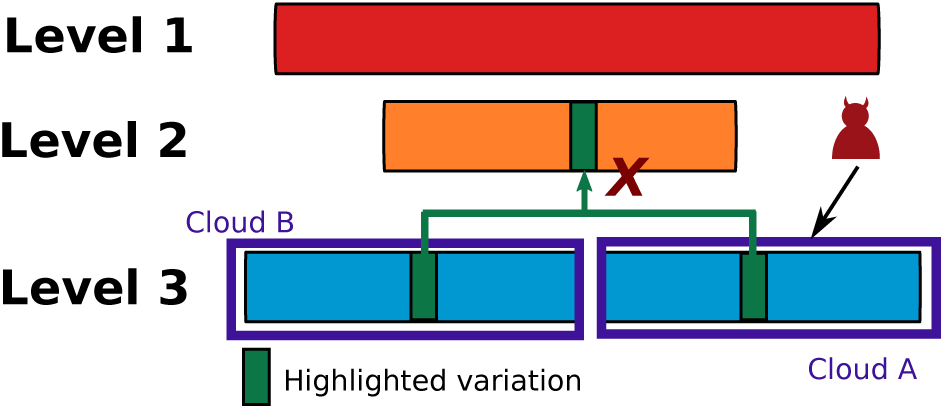
Information stored in two clouds. Partition of the stored data in two clouds make harder the alleles inference. Once an adversary can observe the information in cloud A, he only has access to partial data.

## Conclusion

In this manuscript, we proposed a novel approach to classify the globally sensitive information in genomic data, in multiple, incremental sensitivity levels. These levels are disconnected based on LD relations and prevent attacks amplification by promoting to higher sensitive levels the SNPs that can lead to more sensitive information. We showed that such classification can leverage the complementary characteristics of different alignment algorithms, if selectively applied to subsets of the data reads, guided by such a risk-aware sensitivity classification, taking the best of each algorithm (performance or security). Our approach improves on the state of the art in terms of privacy × performance product, taking into account the computation time and communication cost to the clouds. Furthermore, our approach is suitable for different levels and different algorithms, even as new algorithms appear.

We presented an implementation with multiple filters to efficiently and automatically detect reads that belong to each privacy-sensitivity level, as soon as they are produced. This filtering approach allows adjusting the protection of reads of different levels, with incremental performance gains resulting in an optimized and stable privacy × performance product. We show that the filtering approach can be combined with existing alignment methods (either cleartext, hybrid, cryptographic). In addition, we believe our approach to be timely, and to be a necessary compromise between perfect security and performance, since the growth in genomics data production pushes biocenters to rely on public clouds, and the performance of cryptographic approaches is not satisfactory.

Finally, future work includes evaluating the applicability of the filtering method to the newest sequencing machines that are now able to produce long reads (e.g., containing thousands of bases). Another challenge consists in tolerating more errors in reads, since the newest technologies are not as accurate as the short reads sequencers [54], [55].

## Acknowledgements

This work was supported by the Fonds National de la Recherche Luxembourg (FNR) through PEARL grant FNR/P14/8149128, and by the Fundação para a Ciência e para a Tecnologia (FCT) through funding of the LASIGE Research Unit, ref. UID/CEC/00408/2013.

## References

[1] M. West, G. S. Ginsburg, A. T. Huang, and J. R. Nevins, “Embracing the complexity of genomic data for personalized medicine,” Genome Research, vol. 16, no. 5, pp. 559–566, 2006.

[2] M. Kayser and P. de Knijff, “Improving human forensics through advances in genetics, genomics and molecular biology,” Nature Reviews Genetics, vol. 12, no. 3, pp. 179–192, 2011.

[3] M. L. Metzker, “Sequencing technologies—the next generation,” Nature reviews. Genetics, vol. 11, no. 1, pp. 31–46, 2010.

[4] N. Homer, S. Szelinger, M. Redman, D. Duggan, W. Tembe, J. Muehling, J. V. Pearson, D. A. Stephan, S. F. Nelson, and D. W. Craig, “Resolving individuals contributing trace amounts of dna to highly complex mixtures using high-density snp genotyping microarrays,” PLoS Genet, vol. 4, no. 8, p. e1000167, 2008.

[5] M. Gymrek, A. L. McGuire, D. Golan, E. Halperin, and Y. Erlich, “Identifying personal genomes by surname inference,” Science, vol. 339, no. 6117, pp. 321–324, 2013.

[6] D. R. Nyholt, C.-E. Yu, and P. M. Visscher, “On jim watson’s apoe status: genetic information is hard to hide,” European Journal of Human Genetics, vol. 17, pp. 147–149, 2009.

[7] J. E. Lunshof, R. Chadwick, D. B. Vorhaus, and G. M. Church, “From genetic privacy to open consent,” Nature reviews. Genetics, vol. 9, no. 5, pp. 406–411, 2008.

[8] R. Wang, Y. F. Li, X. Wang, H. Tang, and X. Zhou, “Learning your identity and disease from research papers: information leaks in genome wide association study,” in Proceedings of the 16th ACM conference on Computer and communications security. ACM, 2009, pp. 534–544.

[9] E. Ayday, J. L. Raisaro, U. Hengartner, A. Molyneaux, and J.-P. Hubaux, Privacy-preserving processing of raw genomic data. Springer, 2014.

[10] X. Zhou, B. Peng, Y. F. Li, Y. Chen, H. Tang, and X. Wang, “To release or not to release: evaluating information leaks in aggregate human-genome data,” in European Symposium on Research in Computer Security. Springer, 2011, pp. 607–627.

[11] V. V. Cogo, A. Bessani, F. M. Couto, and P. Verissimo, “A high-throughput method to detect privacy-sensitive human genomic data,” in 14th ACM Workshop on Privacy in the Electronic Society. ACM, 2015, pp. 101–110.

[12] P. E. Verissimo and A. Bessani, “E-biobanking: What have you done to my cell samples?” Security Privacy, vol. 11, no. 6, pp. 62–65, 2013.

[13] A. Bessani, J. Brandt, M. Bux, V. Cogo, L. Dimitrova, J. Dowling, A. Gholami, K. Hakimzadeh, M. Hummel, M. Ismail et al., “Biobankcloud: a platform for the secure storage, sharing, and processing of large biomedical data sets,” in Workshop on Data Management and Analytics for Medicine and Healthcare, 2015.

[14] T. Takai-Igarashi, K. Kinoshita, M. Nagasaki, S. Ogishima, N. Nakamura, S. Nagase, S. Nagaie, T. Saito, F. Nagami, N. Minegishi, Y. Suzuki, K. Suzuki, H. Hashizume, S. Kuriyama, A. Hozawa, N. Yaegashi, S. Kure, G. Tamiya, Y. Kawaguchi, H. Tanaka, and M. Yamamoto, “Security controls in an integrated biobank to protect privacy in data sharing: rationale and study design,” BMC Medical Informatics and Decision Making, vol. 17, no. 1, p. 100, 2017.

[15] S. D. Constable, Y. Tang, S. Wang, X. Jiang, and S. Chapin, “Privacy-preserving gwas analysis on federated genomic datasets,” BMC Medical Informatics and Decision Making, vol. 15, no. 5, p. S2, 2015.

[16] X. Jiang, Y. Zhao, X. Wang, B. Malin, S. Wang, L. Ohno-Machado, and H. Tang, “A community assessment of privacy preserving techniques for human genomes,” BMC Medical Informatics and Decision Making, vol. 14, no. 1, p. S1, 2014.

[17] P. R. Payne, N. H. Shah, J. D. Tenenbaum, and L. Mangravite, Democratizing Health Data for Translational Research. WORLD SCIENTIFIC, 2017, pp. 240–246.

[18] A. Ardeshirdavani, E. Souche, L. Dehaspe, J. Van Houdt, J. R. Vermeesch, and Y. Moreau, “Ngs-logistics: federated analysis of ngs sequence variants across multiple locations,” Genome Medicine, vol. 6, no. 9, p. 71, 2014.

[19] T. G. A. for Genomics and Health, “A federated ecosystem for sharing genomic, clinical data,” Science, vol. 352, no. 6291, pp. 1278–1280, 2016.

[20] S. O. Dyke, E. S. Dove, and B. M. Knoppers, “Sharing health-related data: a privacy test?” npjgenmed, vol. 1, p. 16024, 2016.

[21] Y. Li, C. J. Willer, J. Ding, P. Scheet, and G. R. Abecasis, “Mach: using sequence and genotype data to estimate haplotypes and unobserved genotypes,” Genetic epidemiology, vol. 34, no. 8, pp. 816–834, 2010.

[22] K. B. Jacobs, M. Yeager, S. Wacholder, D. Craig, P. Kraft, D. J. Hunter, J. Paschal, T. A. Manolio, M. Tucker, R. N. Hoover et al., “A new statistic and its power to infer membership in a genome-wide association study using genotype frequencies,” Nature genetics, vol. 41, no. 11, pp. 1253–1257, 2009.

[23] A. Michalas, N. Paladi, and C. Gehrmann, “Security aspects of e-health systems migration to the cloud,” in e-Health Networking, Applications and Services (Healthcom), 2014 IEEE 16th International Conference on. IEEE, 2014, pp. 212–218.

[24] B. Fabian, T. Ermakova, and P. Junghanns, “Collaborative and secure sharing of healthcare data in multi-clouds,” Information Systems, vol. 48, pp. 132–150, 2015.

[25] Y. Tong, J. Sun, S. S. Chow, and P. Li, “Cloud-assisted mobile-access of health data with privacy and auditability,” IEEE Journal of biomedical and health Informatics, vol. 18, no. 2, pp. 419–429, 2014.

[26] B. Yüksel, A. Küpçü, and Ö. Ö zkasap, “Research issues for privacy and security of electronic health services,” Future Generation Computer Systems, vol. 68, pp. 1–13, 2017.

[27] M. T. Goodrich, “The mastermind attack on genomic data,” in Security and Privacy. IEEE, 2009, pp. 204–218.

[28] K. Kidd, A. Pakstis, W. Speed, and E. e. a. Grigorenko, “Developing a snp panel for forensic identification of individuals,” Forensic science international, vol. 164, no. 1, pp. 20–32, 2006.

[29] B. Malin, Compromising privacy with trail re-identification: the REIDIT algorithms. Carnegie Mellon University. Center for Automated Learning and Discovery, 2002.

[30] B. Malin, Protecting dna sequence anonymity with generalization lattices. Carnegie Mellon University, School of Computer Science [Institute for Software Research International], 2004.

[31] B. Malin and L. Sweeney, “How (not) to protect genomic data privacy in a distributed network: using trail re-identification to evaluate and design anonymity protection systems,” Journal of biomedical informatics, vol. 37, no. 3, pp. 179–192, 2004.

[32] L. T. Vaszar, M. K. Cho, and T. A. Raffin, “Privacy issues in personalized medicine,” Pharmacogenomics, vol. 4, no. 2, pp. 107–112, 2003.

[33] R. B. Altman and T. E. Klein, “Challenges for biomedical informatics and pharmacogenomics,” Annual review of pharmacology and toxicology, vol. 42, no. 1, pp. 113–133, 2002.

[34] M. Naveed, E. Ayday, E. W. Clayton, J. Fellay, C. A. Gunter, J.-P. Hubaux, B. A. Malin, and X. Wang, “Privacy in the genomic era,” ACM Computing Surveys (CSUR), vol. 48, no. 1, p. 6, 2015.

[35] B. Malin and L. Sweeney, “Determining the identifiability of dna database entries.” in Proceedings of the AMIA Symposium. American Medical Informatics Association, 2000, p. 537.

[36] Z. Lin, M. Hewett, and R. B. Altman, “Using binning to maintain confidentiality of medical data.” in Proceedings of the AMIA Symposium. American Medical Informatics Association, 2002, p. 454.

[37] M. Kantarcioglu, W. Jiang, Y. Liu, and B. Malin, “A cryptographic approach to securely share and query genomic sequences,” IEEE Transactions on information technology in biomedicine, vol. 12, no. 5, pp. 606–617, 2008.

[38] T. Neubauer and J. Heurix, “A methodology for the pseudonymization of medical data,” International Journal of Medical Informatics, vol. 80, no. 3, pp. 190–204, 2011.

[39] B. A. Malin, “An evaluation of the current state of genomic data privacy protection technology and a roadmap for the future,” Journal of the American Medical Informatics Association, vol. 12, no. 1, pp. 28–34, 2005.

[40] M. Humbert, E. Ayday, J.-P. Hubaux, and A. Telenti, “Addressing the concerns of the lacks family: quantification of kin genomic privacy,” in Proceedings of the 2013 ACM SIGSAC conference on Computer & communications security. ACM, 2013, pp. 1141–1152.

[41] M. C. Schatz, “Cloudburst: highly sensitive read mapping with mapreduce,” Bioinformatics, vol. 25, no. 11, pp. 1363–1369, 2009.

[42] R. V. Pandey and C. Schlötterer, “Distmap: a toolkit for distributed short read mapping on a hadoop cluster,” PLoS One, vol. 8, no. 8, pp. 1363–1369, 2013.

[43] Y. Huang, D. Evans, J. Katz, and L. Malka, “Faster secure two-party computation using garbled circuits.” in USENIX Security Symposium, vol. 201, no. 1, 2011.

[44] E. De Cristofaro, S. Faber, and G. Tsudik, “Secure genomic testing with size-and position-hiding private substring matching,” in Proc. of the 12th ACM Workshop on Privacy in the Electronic Society, 2013, pp. 107–118.

[45] Y. Chen, B. Peng, X. Wang, and H. Tang, “Large-scale privacy-preserving mapping of human genomic sequences on hybrid clouds.” in NDSS, 2012.

[46] V. Popic and S. Batzoglou, “Privacy-preserving read mapping using locality sensitive hashing and secure kmer voting,” bioRxiv, 2016.

[47] B. H. Bloom, “Space/time trade-offs in hash coding with allowable errors,” Commun. ACM, vol. 13, no. 7, pp. 422–426, 1970.

[48] R. Schnell, T. Bachteler, and J. Reiher, “Privacy-preserving record linkage using bloom filters,” BMC Medical Informatics and Decision Making, vol. 9, no. 1, p. 41, 2009.

[49] “1000 Genomes Project: A Deep Catalog of Human Genetic Variation,” available at: http://www.1000genomes.org/.

[50] S. Sankararaman, G. Obozinski, M. I. Jordan, and E. Halperin, “Genomic privacy and limits of individual detection in a pool,” Nature genetics, vol. 41, no. 9, pp. 965–967, 2009.

[51] F. Takeuchi, K. Yanai, T. Morii, Y. Ishinaga, K. Taniguchi-Yanai, S. Nagano, and N. Kato, “Linkage disequilibrium grouping of single nucleotide polymorphisms (snps) reflecting haplotype phylogeny for efficient selection of tag snps,” Genetics, vol. 170, pp. 291–304, 2005.

[52] K. G. Ardlie, L. Kruglyak, and M. Seielstad, “Patterns of linkage disequilibrium in the human genome,” Nature Review Genetics, vol. 3, pp. 299–309, 2002.

[53] J. Baron, K. El Defrawy, K. Minkovich, R. Ostrovsky, and E. Tressler, “5pm: Secure pattern matching,” in Security and Cryptography for Networks. Springer, 2012, pp. 222–240.

[54] T. C. Glenn, “Field guide to next-generation dna sequencers,” Molecular ecology resources, vol. 11, pp. 759–769, 2011.

[55] C. Bleidorn, “Third generation sequencing: technology and its potential impact on evolutionary biodiversity research,” Systematics and Biodiversity, vol. 14, pp. 1–8, 2016.

